# Enhancing plant photosynthesis through activating cells deficient in photosynthetic apparatus

**DOI:** 10.1101/2023.09.27.559740

**Authors:** Fenfen Miao, Yong-yao Zhao, Mingju Lyu, Bihai Shi, Fusang Liu, Xin-Guang Zhu

## Abstract

Much efforts have been devoted to identify options to improve plant photosynthetic capacity. Most of these engineering focuses more on enhancing photosynthetic capacity of the mesophyll cells. In this study, we specifically expressed *CYTOKININRESPONSIVE GATA FACTOR 1 (CGA1)* in the vascular bundle and bundle sheath cells (*Pro_GLDPA_: CGA1*) which generally have low photosynthetic capacity. The introduction of *CGA1* resulted in increased numbers and sizes of chloroplasts in both bundle sheath and vascular bundle, without noticeable difference in these properties in mesophylls. Additionally, compared with wild type (WT), transgenic lines showed enhanced development of mitochondria and peroxisomes. Leaf photosynthetic rates in these transgenic lines were higher than those in WT, especially under high light. Furthermore, the amount of RuBisCO, V_cmax_, J_max_, and the number of starch granules were also significantly increased in the transgenic lines, which led to about 20% increase in biomass. Remarkably, the CO_2_ compensation point of *Pro_GLDPA_: CGA1* decreased compared with WT, resembling the characteristics of proto-Kranz during C_4_ evolution. All these results indicated that enhancing photosynthetic properties of vascular bundle and bundle sheath cell is an effective approach to improve leaf photosynthesis, and highlighted that utilization of the cell lacking photosynthetic capacity is a promising way to improve plant photosynthesis for greater capacity.

## Introduction

Improving photosynthesis is an effective approach to increase crop yield potential (South et al. 2019; Zhu et al. 2010); it is also a potential approach to mitigate global change (IPCC 2007; Goli et al. 2016). Much effort has been devoted to increasing photosynthetic capacity (Tanaka et al. 2013; South et al. 2019; Chen et al. 2020; De Souza et al. 2022; Wei et al. 2022; Zhu et al. 2010). For C_3_ plants, mesophyll cell (MC) is the major cell type for photosynthesis (Lei Hua 2021; Sylvain Aubry 2014). Most current efforts of improving photosynthetic capacity in C_3_ plants are related to manipulating photosynthetic machinery in MC (South et al. 2019; De Souza et al. 2022; Shen et al. 2019). Compared with MC, many other cell types have deficient photosynthetic capacity (Sylvain Aubry 2014; Lei Hua 2021; Simkin and 2020; Hu et al. 2019). The potential to gain increased plant photosynthetic productivity through enhancing photosynthetic capacity of these cells has not been studied.

Chloroplast is an organelle that carry out photosynthesis. *GATA NITRATE-INDUCIBLE CARBON-METABOLISM-INVOLVED (GNC) and GOLDEN TWO-LIKE (GLK)* family transcription factors control the biogenesis, development and division of chloroplasts (Yi-Hsuan Chiang 2012; Zubo et al. 2018; Cackett et al. 2022; Dong-Yeon Lee 2021; Wang et al. 2017), and they can positively regulate and coordinate the expression of photosynthesis related genes(Raines et al. 2020; Liu et al. 2018; Waters et al. 2009; Darryl Hudson 2013). Therefore, these transcription factors have the potential to be used as molecular tools to boost photosynthetic capacities of photosynthetic inactive tissues. Indeed, over-expression of *GLK* promotes the chloroplast development and results in photosynthetic activity in roots and calli (Koichi Kobayashi and Masuda 2013; Nakamura et al. 2009). In addition, when the GLK or GNC family transcription factors are constitutively expressed in rice regardless of cell types or controlled by their own promoter, leaf photosynthesis and biomass are both increased (Lee et al. 2021; Yeh et al. 2022; Li et al. 2020). However, when *CYTOKININ-RESPONSIVE GATA1* (*CGA1, the member of GNC)* or *GLK* is specifically expressed in rice bundle sheath cells, the photosynthetic performance of leaf did not show apparent change (Wang et al. 2017; Dong-Yeon Lee 2021). Though these different transcription factors have the power to boost chloroplast development, whether cell-specific expression of these transcription factors in cells with deficient photosynthetic capacity can result in significantly enhanced leaf photosynthetic capacity still needs to be tested.

The main function of vascular tissue include transport of water, ions, photoassimilates and other nutrients (Hardtke 2023). Bundle sheath cells, which surround vascular bundle, are specialized in the water transport and sulfur transport in rice (Lei Hua 2021), or glucosinolate biosynthesis and sulfur in Arabidopsis (Sylvain Aubry 2014). Earlier studies show that cells around the vascular bundle do have photosynthetic activity (Hibberd and Quick 2002; Gao et al. 2018), though the photosynthetic capacity of these vascular related tissues including bundle sheath cells is generally weaker in C_3_ plants (Lei Hua 2021; Hunt et al. 2023; Kim et al. 2021; Sylvain Aubry 2014; Pyke* 1998), and the potential of their contribution to whole leaves or plants photosynthesis is largely unclear. In the case of C_4_ species, bundle sheath cells play a crucial role in the photosynthesis (Langdale 2011; Sage 2012).

In this study, we used the promoter of the *glycine decarboxylase P-protein (GLDP)* in *Flaveria trinervia*, which guides expression of genes in the vascular bundle cell (VBC) and bundle sheath cell (BSC) in *Arabidopsis* (Engelmann et al. 2008), to drive the expression of *CGA*1 in VBC and BSC in *Arabidopsis*. This study aims to test the hypothesis that the increased photosynthetic capacity of vascular bundle cell (VBC) and bundle sheath cell (BSC) can substantially improve the leaf photosynthesis of *Arabidopsis*.

## Method

### Plant materials and growth conditions

*Arabidopisis* plants used here are of the ecotype Columbia (Col-0). Seeds of wild-type and transgenic lines *(Pro_GLDPA_-CGA1-GFP-L3/L4/L9*) were stratified at 4°C for 3 days and then sown into peat soils. The wild-type and transgenic lines were grown in a growth chamber (gREEN FUTURE) under short day conditions, with 10 h of light (150 μmol m^−2^s^−1^) and 14 h of darkness, with 60% relative humidity under 21±1°C. The morphology for leaves and whole plant of wild type and transgenic lines at the 30 days were photographed.

### Plasmid construction and plant transformation

The *GLYCINE DECARBOXLASE P-SUBUNIT A* (*GLDPA*) from *Flaveria trinervia* was amplified from genomic DNA by Polymerase Chain Reaction (PCR) with the primer pair Pro_GLDPA_-F and Pro_GLDPA_-R (supplementary table 1). After sequence verification, the resulting fragment was inserted into *EcoRI* cloning sites of the pCAMBIA1300 expression cassette with hygromycin resistance, yielding the construct *Pro_GLDPA_-pCAMBIA1300*. Next, the *Arabidopsis* gene CGA1 (AT4G26150) was amplified from the cDNA with the primer pair CGA1-F and CGA1-R along with the Green Fluorescent Protein (GFP) coding sequence. Then the resulting PCR fragment of CGA1 and GFP as template and CGA1-F and GFP-R as primers yielded fusion PCR fragment CGA1-GFP. Last, CGA1-GFP was inserted into *SacI* cloning sites in *Pro_GLDPA_-pCAMBIA1300*, yielding the *Pro_GLDPA_-CGA1-GFP-pCAMBIA1300* construct for plant transformation.

To verify GLDPA specific expression in *Arabidopsis*, we first cloned the fragment of GLDPA promoter, Pro_GLDPA_. Specifically, the primer pair 1381-Pro_GLDPA_-F and 1381-Pro_GLDPA_-R were used to amplify the fragment in Pro_GLDPA_, which was then inserted into the *GUS-*pCAMBIA1381 construct with hygromycin resistance to yield *Pro_GLDPA_-GUS-pCAMBIA1381* construct. The primer Venus-Pro_GLDPA_-F and Venus-Pro_GLDPA_-R were also used to amplify the fragment of Pro_GLDPA_, which generated *Pro_GLDPA_-Venus-N7* construct with knnamycin resistance. All amplified fragments were confirmed by sequencing. The sequences of primers used for plasmid constructions are listed in Supplementary Table 1.

Transgenic Arabidopsis plants were generated by a floral dip method (Clough and Bent 1998) with the Agrobacterium tumefaciens strain GV3101. First, transgenic seeds were surface sterilized with ethanol. Then, according to their corresponding resistance, seeds were plated on half-strength Murashige and Skoog (MS) medium (0.8% agar) containing 25mg l^−1^ hygromycin or 80 mg l^−1^ knnamycin. They were then stratified in the dark at 4°C for 3 days, then transferred to long-day conditions, with 16 h of light (100 μmol m^−2^s^−1^) and 8h of darkness, with 60% relative air humidity at 22°C. Homozygous transgenic (T2) plants were screened and confirmed by seed germination on the corresponding antibiotic-containing medium in combination with PCR analyses (Chen et al. 2020). The sequences of primers for transgenic identification are listed in Supplementary Table1.

### RNA extraction and RT-qPCR

The RT–qPCR analysis was conducted as described previously (Chen et al. 2020). Briefly, the illuminated new fully expanded leaf of 28-day-old plants were used to isolate total RNA with the PureLink RNA Mini Kit (Life Technologies Corporation, USA) according to manufacturer’s protocol. The isolated total RNA (2μg) was used as templates to produce the cDNA through TranScript One-Step gDNA Removal and cDNA Synthesis SuperMix (TransGen Biotech, China). Real-time qPCR was performed with SYBR Green Real-Time PCR Master Mix (Yeasen, China) using the CFX 96 system (Bio-Rad). The relative transcript levels were determined according to the comparative threshold cycle method (Livak and Schmittgen 2001). AtACTIN8 (AT3G18780) was used as an internal control to analyze the expression level. The sequences of primers are listed in Supplementary Table 1.

### Transmission electron microscopy (TEM) imaging

For TEM observation, about 1 mm sections of new fully expanded leaf at 28-day-old were harvested from WT and Line3, Line4 and Line9 at two o’clock in the afternoon. Then, the samples were dehydrated, and embedded following the standard protocol (Wang et al. 2020). Ultrathin sections of the samples were cut with a diamond knife on a Leica EMUC6-FC6 ultramicrotome, and then imaged at 80 kV with a Hitachi H-7650 transmission electron microscope.

Numbers and sizes of planar cell area, chloroplast, mitochondria, peroxisome and starch were quantified by ImageJ software (Schneider et al. 2012) with TEM images of *Arabidopsis* leaves from WT and transgenic lines. Plasmodesmata between bundle sheath cell and mesophyll cell (BSC-MC), bundle sheath cell and bundle sheath cell (BSC-BSC), bundle sheath cell and vein (BSC-Vein) were analyzed using the TEM images.

### TEM for quantifying immunogold labeling

The sampling procedure used here was same as that for the TEM assay. The leaf segments were detached from WT, Line 3, Line 4 and Line 9 plants, and fixed in 4% paraformaldehyde (PFA) and 0.1% GA, stayed in 4% PFA and 0.1% GA for 4 hours, then transferred into 2% PFA at 4□ overnight. The fixed samples were washed by phosphate buffer, followed by 0.1 M glycine in PBS buffer for 1 h, then dehydrated with gradient 30%, 50%, and 70% ethanol at 4□, and then at 85%, 95% and 100% ethanol at −20□. LR-white resin was used (Ted Pella, Redding, CA) for infiltration and embedding, blocks were polymerized under UV light at −10□ for 3 days. 80 nm ultra-thin sections were cut by ultramicrotome (UC7, Leica Microsystem) and collected on formvar supported nickel single slot grids, which were floated on PBS for 15 min and then on PBS and 1% bovine serum albumin for 15 min. The sections were incubated with the primary antibody (dilution in 0.1 M PBS and 1% bovine serum albumin, Rubisco, 1:1600; GDCP, 1:800, v/v) at 4□ overnight, washed by PBS and 0.1% BSA for 4 times, then treated with secondary antibody (Anti-rabbit IgG, 10 nm gold particle, dilution at 1:50 v/v, G7402, Sigma) at room temperature for 1h. The grids were post-stained with 3% uranyl acetate, and observed under a transmission electron microscopy (Talos L120C, Thermo Fisher) at 120 kV and imaged by a Ceta2 camera. The density of gold labelling was counted and calculated by ImageJ software.

### Gas exchange measurements

Gas exchange measurements were conducted using a LI-COR 6800XT portable photosynthesis system (LI-COR Biosciences, Lincoln, USA). The mature expanded leaves of 28-day-old plants were used. Measurements were performed under a photosynthetic photon flux density (PPFD) of 150 μmol m^−2^ s^−1^ and a CO_2_ concentration of 400 ppm. To record the light response curve of photosynthetic CO_2_ uptake rate (*A*), the CO_2_ concentration was set to 400 ppm, and PPFD was stetted as 1200, 800, 600, 400, 300, 200, 150, 100, 50, and 0 µmol m^−2^ s ^−1^. For the CO_2_-response curve (A-Q curve), the PPFD was set to 600 µmol m^−2^ s ^−1^, and the CO_2_ concentration was gradually decreased from 400 to 30 ppm, which include 400, 250, 150, 100, 80, 50, 30 ppm, and then increased from 400 to 700, 1000, 1300, 1500, and 1800 ppm. CO_2_ compensation point was determined using linear regression of *A* and Ci measurements under ambient CO_2_ levels of 30, 50, 70, 90,150 ppm under a PPFD of 250 µmol m^−2^s^−1^ (Sharkey 1988). A-Q curves were fitted by a non-rectangular hyperbola model (Eqn. 1). *A(I)* is *A* under a particular incident light intensity(*I*). *A_sat_* is leaf photosynthetic rate under saturating incident light intensity. ¢is the quantum yield of CO_2_. 0 represents the convexity of the A-Q curve, *R_d_* is day respiration.

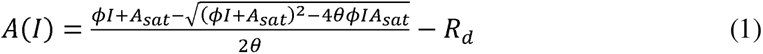

A-Ci curves were fitted based on Farquhar-von Caemmerer-Berry (FvCB) model (Farquhar et al. 1980) using the R package ‘plantecophys’ (Duursma 2015). The RuBP saturated maximal photosynthetic electron transport rate (Jmax) and The RuBP and CO_2_ saturated maximal rate of ribulose1,5-bisphosphate (RuBP) carboxylation by Rubisco (Vcmax) and R_d_ were estimated by the fitted FvCB model.

### High light intensity treatment

High light acclimation was performed as described previously with modifications (Ma et al. 2014). Specifically, 21-day-old *Arabidopsis* plants were transferred from 200 μmol m^−2^s ^−1^ (normal light intensity, NL) to 250 μmol m^−2^ s ^−1^ for 2 days, then exposed to 300 μmol m^−2^ s ^−1^ for 3 days, then exposed to 450 μmol m^−2^ s ^−1^ for 4 days (high light intensity, HL). After high light intensity treatment, leaf number and fresh weight were recorded. Phenotypes of wild type and the transgenic lines of *Arabidopsis* grown for 30-day-old under high light intensity were photographed.

### Protein content measurement

The protein content measurement was conducted as described previously with some modifications (Wei et al. 2022). Briefly, new fully expanded leaves under illumination were sampled from 28-day-old *Arabidopsis* plants. For each sample, the leaf area of 0.84 cm^2^ was sampled and grounded in liquid nitrogen and homogenized with 200 μL extraction buffer (20 mM Tris (pH 7.5), 100 mM NaCl, 2.5 mM MgCl2, 1 mM EGTA, 1 mM DTT and protease inhibitor cocktail (1:50) (Roche). After gentle shaking on ice for 30 min, the homogenates were centrifuged at 12,000 g for 20 min at 4 °C. The supernatants were used for protein content quantification by the Quick Start Bradford Reagent (Bio-rad).

### Measurement of chlorophyll content

Leaf discs (0.84 cm^2^) were taken from fully expanded leaves, and chlorophyll was extracted with 80% acetone at 4 °C for 24 h in darkness, then the supernatant was used for the absorbance measurement at 652 nm in a spectrophotometer (Arnon 1949). The total chlorophyll content was calculated with the following formula:

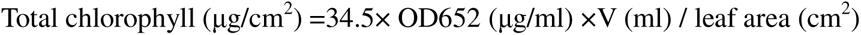

### Measurement of biomass and leaf number

The aboveground part of *Arabidopsis* was cut from a 30-day-old plants and weighed to get the fresh weight. Leaf number were counted and recorded.

### Confocal microscopy

The leaves were detached from WT and *Pro_GLDPA_-Venus-N7* transgenic lines, and then fixed with 4% (w/v) PFA for 120 min in PBS under vacuum at room temperature. The ClearSee assays were performed as described previously (Kurihara et al. 2015), and the fixed tissues were cleared with ClearSee solution at room temperature for 5 days until the leaves fully decolorized. ClearSee solutions were prepared by mixing xylitol powder (10% (w/v)), sodium deoxycholate (15% (w/v)) and urea (25% (w/v)] in H_2_O. ClearSee-treated leaves were observed by confocal microscopy. Fluorescence signals were recorded under a Zeiss LSM 880 confocal laser scanning microscope. To observe the Venus signals, an excitation wavelength of 488□nm from an argon laser was used, and the emission signal was detected at 500–550□nm.

### β-Glucuronidase (GUS) reporter assay

The leaves of Pro_GLDPA_-GUS were placed in β-glucuronidase (GUS) staining solution (0.1M Na_2_HPO_4_, 0.1M NaH_2_PO_4_, 5 mM K_3_[Fe (CN)_6_], 5 mM K_4_[Fe (CN)_6_], 1% Triton X −100, 20% methanol, 0.6 mg/ml 5-bromo-4-chloro-3-indolyl-beta-D-glucuronic acid, cyclohexylammonium salt (X-Gluc)). After incubation at 37°C under dark overnight, the tissue was destained by washing several times with 70% (v/v) ethanol. Nonsectioned leaf was imaged under a Leica Stereomicroscope MZ12.5.

### SDS-page and western blot

Same leaf discs (0.84 cm^2^) were taken from fully expanded leaves of 28 days WT and line3, line4, and line9 plants. Leaf-soluble proteins were extracted by grinding the frozen leaves in 200 μL extraction buffer containing 20 mM Tris (pH 7.5), 100 mM NaCl, 2.5 mM MgCl2, 1 mM EGTA, and 1 mM DTT. After denaturation, the debris was removed by centrifugation at 12000 g for 10 min at 4 °C. The same leaf area (0.84 cm^2^) is used for quantification, and samples of 10 μL protein were loaded onto each lane. The gels were stained with brilliant blue R-250, de-stained with 20% methanol, and then observed.

For western blot analysis, the extraction of total protein following the same procedure as for sodium dodecyl sulfate–polyacrylamide gel electrophoresis (SDS-PAGE) experiment. Samples of 10 μL extraction were separated by SDS-PAGE and transferred to PVDF membranes (Amersham, GE Healthcare). After blocking with 5% skim milk in TBST buffer (20 mM Tris/HCl, pH 7.6, 137 mM NaCl, 0.1% Tween) at 4 °C overnight, membranes were incubated with primary antibodies raised against GFP (Abmart, M20004, 1:2000) at the dilutions recommended by the supplier, followed by incubation with secondary antibody (goat anti-rabbit, Abclone) at a dilution of 1: 1:5000. Membranes were imaged using a chemiluminescence imager device (Tanon-5200) after incubation in BeyoECL Plus (beyotime, P0018S) for 5 min at room temperature.

### Statistical analysis and accession number

The graph in this study was plotted by Graphpad prism version 8. Mann-Whitney U test and student’s t-test (unpair, two-side) in this study was both performed by Graphpad prism version 8. The gene accession number of *CYTOKININ-RESPONSIVE GATA1* (*CGA1)* is AT4G26150.

## Result

### Expressing *CGA1* in the vascular bundle (VB) and bundle sheath (BS) resulted in greener veins and increased chlorophyll content

We linked the promoter of *Flaveria trinervia GLDPA* to *GUS* and *Venus* tags separately (Supplementary Fig 1A). The signals of GUS (Supplemental Fig 1B, C) and Venus (Supplemental Fig 1D, E) were only observed in the vascular tissue, which indicated that the promoter of *Flaveria trinervia GLDPA* can tissue-specific drive gene expression in the vascular tissue of *Arabidopsis*, as reported earlier (Engelmann et al. 2008). *CYTOKININRESPONSIVE GATA FACTOR 1 (CGA1)* expressed in both C_3_ and C_4_ leaf, and CGA1 mainly expressed in leaf of *Arabidopsis* (Supplementay fig 2). We generated three *CGA1-GFP* transgenic lines of *Arabidopsis* under control of *Flaveria trinervia GLDPA* promoter (Transgenic plant named *Pro_GLDPA_: CGA1*, three lines are L3, L4, L9, respectively) (Fig 1A). As expected, compared with wild-type (WT), the large increase in the transcription levels of *CGA1* was detected in the transgenic lines (Fig 1B). The chlorophyll concentrations in the transgenic lines increased significantly as well (Fig 1C), consistent with the greener veins on both the abaxial and the adaxial sides (Fig 1D).

**Fig 1.**
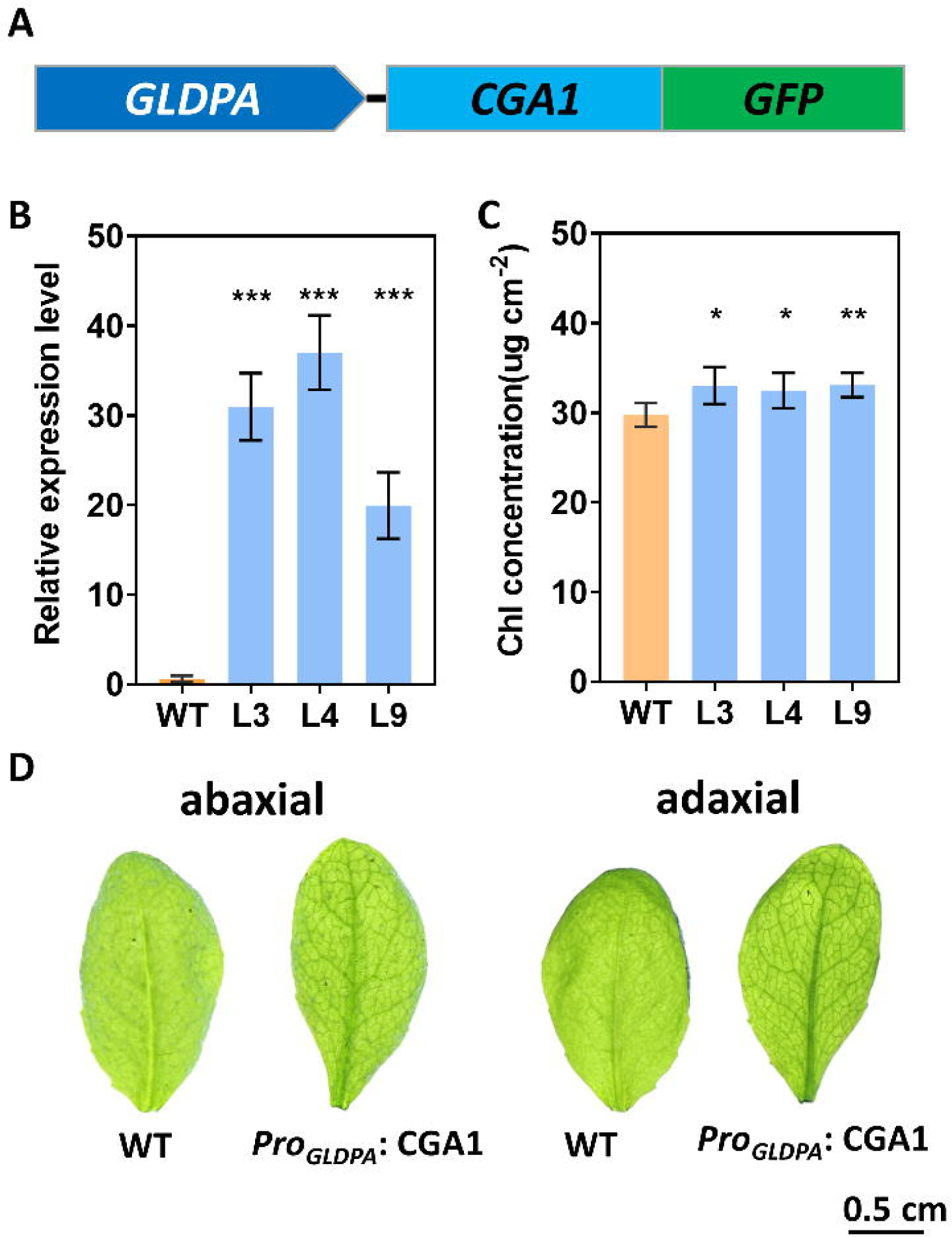
Expression of *CGA1* in the bundle sheath and vascular bundle resulted in greener veins and increased chlorophyll content. (A) Schematic presentation of the *Pro_GLDPA_*-CGA1-GFP that was transformed into *Arabidopsis* to express *CGA1* in the vascular tissue. (B) Expression levels of *CGA1* in leaves of wild type (WT) and transgenic lines of *Arabidopsis Pro_GLDPA_*: CGA1 (L3/L4/L9), which were analysed by RT–qPCR (n=3). The specific primers (CGA1-F and CGA1-R), were used to examine the expression level of *CGA1* in leaves. Bars indicate the SD, \**P*<0. 5,\*\**P*<0.01,\*\*\**P*<0.001, two-sided student’s T-test, unpaired. (C) Chlorophyll concentration for the WT and transgenic lines (n=5). Bars indicate the SD, \**P*<0. 5,\*\**P*<0.01,\*\*\**P*<0.001, two-sided student’s T-test,unpaired. (D) Photographs of the leaf blades of WT and transgenic lines (*Pro_GLDPA_*: CGA1). Scale bars, 0.5 cm.

### Expressing *CGA1* resulted in greater chloroplast number and size in bundle sheath and vascular bundle

To further test whether *CGA1* could promote the development of chloroplasts in bundle sheath cells (BSC) and vascular bundle cells (VBC), ultrathin sections were performed to examine the anatomy of chloroplast. As shown in Fig 2A-F (WT: A, C, E; *Pro_GLDPA_: CGA1*: B, D, F), the numbers and sizes of chloroplasts in the VBC and BSC in transgenic lines were significantly increased compared with WT (Fig 2C, D, E, F, I, J, K, L, M). The numbers of chloroplasts in VBC and BSC in transgenic lines were twice as many as those in the WT (Fig 2I, M). The sizes of chloroplasts in transgenic plants were about 1.4 times of that for WT (Fig 2J). We also found that the area of BSC (Fig 2G), the total chloroplast area in per BSC (Fig2K) and the ratio of chloroplast area in BSC area were all significantly increased (Fig 2L). These results indicated that the expression of *CGA1* in the bundle sheath and vascular bundle promotes chloroplast biogenesis and division. There was no significant difference in the number of chloroplasts per MC and MC area between transgenic line and WT (Supplementary Fig 3A, B), which indicated that the expression of *CGA1* did not influence MC. Furthermore, the starch granule numbers in the BSC and VBC of transgenic lines were dramatically increased by about 2-5 times (Fig 2H, N).

**Fig 2.**
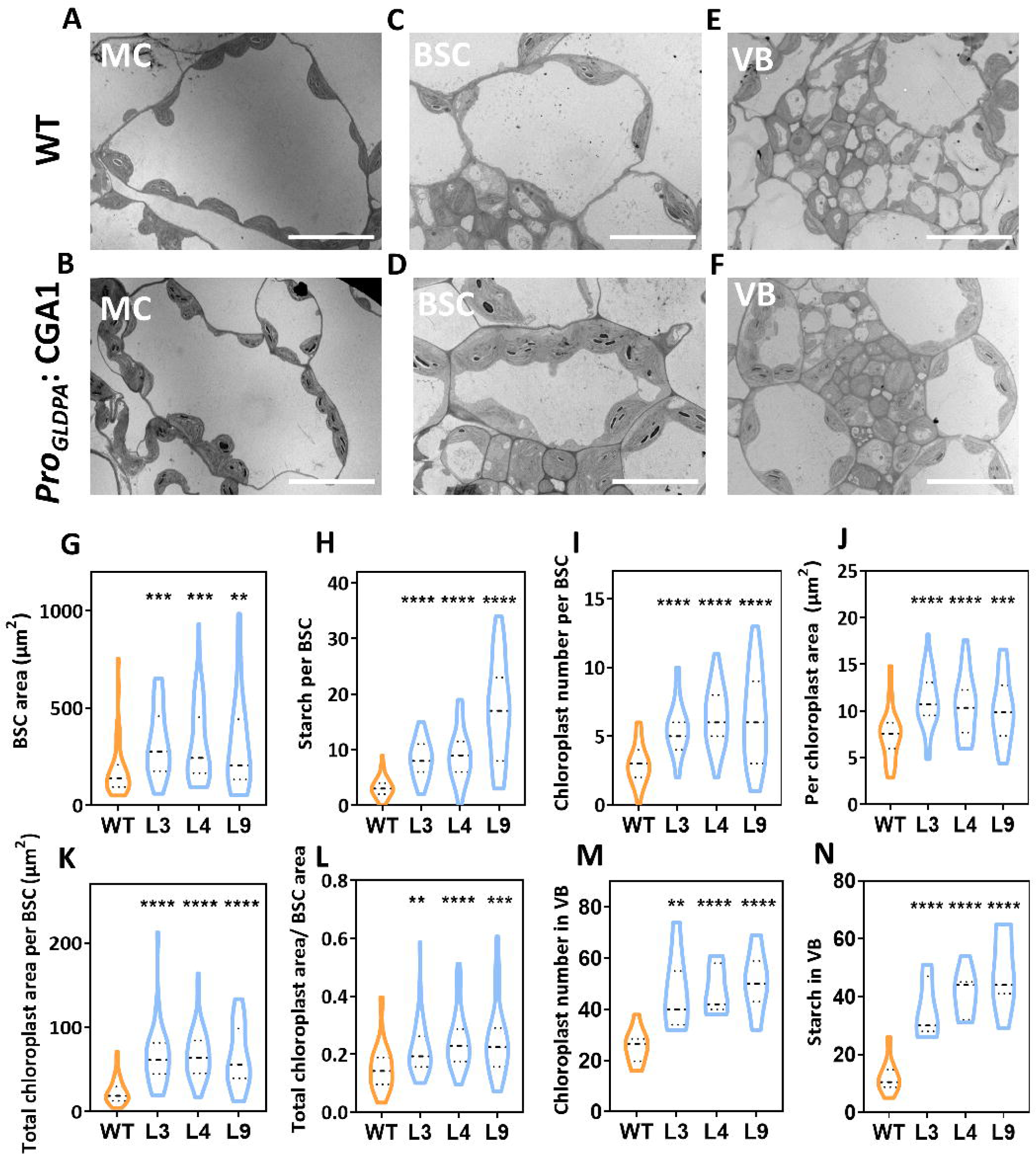
Expression of *CGA1* in the bundle sheath and vascular bundle increased chloroplast number and size in bundle sheath cell and vascular bundle cell. (A to F) Transmission electron micrographs of mesophyll cell (MC), bundle sheath cell (BSC) and vascular bundle (VB). Wild-type (WT) (A, C, E) and transgenic lines of Arabidopsis *Pro_GLDPA_:* CGA1 (B, D, F). Scale bars, 20 μm (A and B) and 10 μm (C–F). (G-N) Violine plot showing BSC area, and chloroplast size and number in BSC and VB. (G) The average size of each BSC (n ≥39). (H) The starch granule number in each BSC (n ≥39). (I) The average chloroplast number in each BSC (n ≥39). (J) The average size of each chloroplast in each BSC (n ≥39). (K) The average total chloroplast area in each BSC (n ≥39). (L) The ratio of chloroplast area in BSC (n ≥39). (n ≥ 39 cells and from 3 individuals representing at least three independent lines in each case). (M) The average chloroplast number in VB (n ≥7). (N) The average starch granule number in VB (n ≥7).). (n ≥ 7 VB and from 3 individuals representing at least three independent lines in each case) \*\**P*<0.01\*\*\**P*<0.001, \*\*\*\**P*<0.0001, two-sided student’s *t*-test,unpaired.

We further examined the expression of RuBisCO using Immunogold labelling technique. As shown in the picture, the density of immunogold labeled RuBisCO in BSC in transgenic lines was higher than that in the WT (Fig 3A, B). The total number of immunogold particles per BSC increased by about 1 fold and the density of RuBisCO in each BSC area (um^2^) was also significantly increased (Table 1). There was no significant difference in the density of immunogold particles on planar chloroplast area (um^2^) between transgenic line and WT (Table 1), suggesting the increase abundance of RuBisCO was due to increase of the number and size of chloroplast. The increased starch granules and RuBisCO content further implied that the increased chloroplasts in the VB and BS are photosynthetically active.

**Fig 3.**
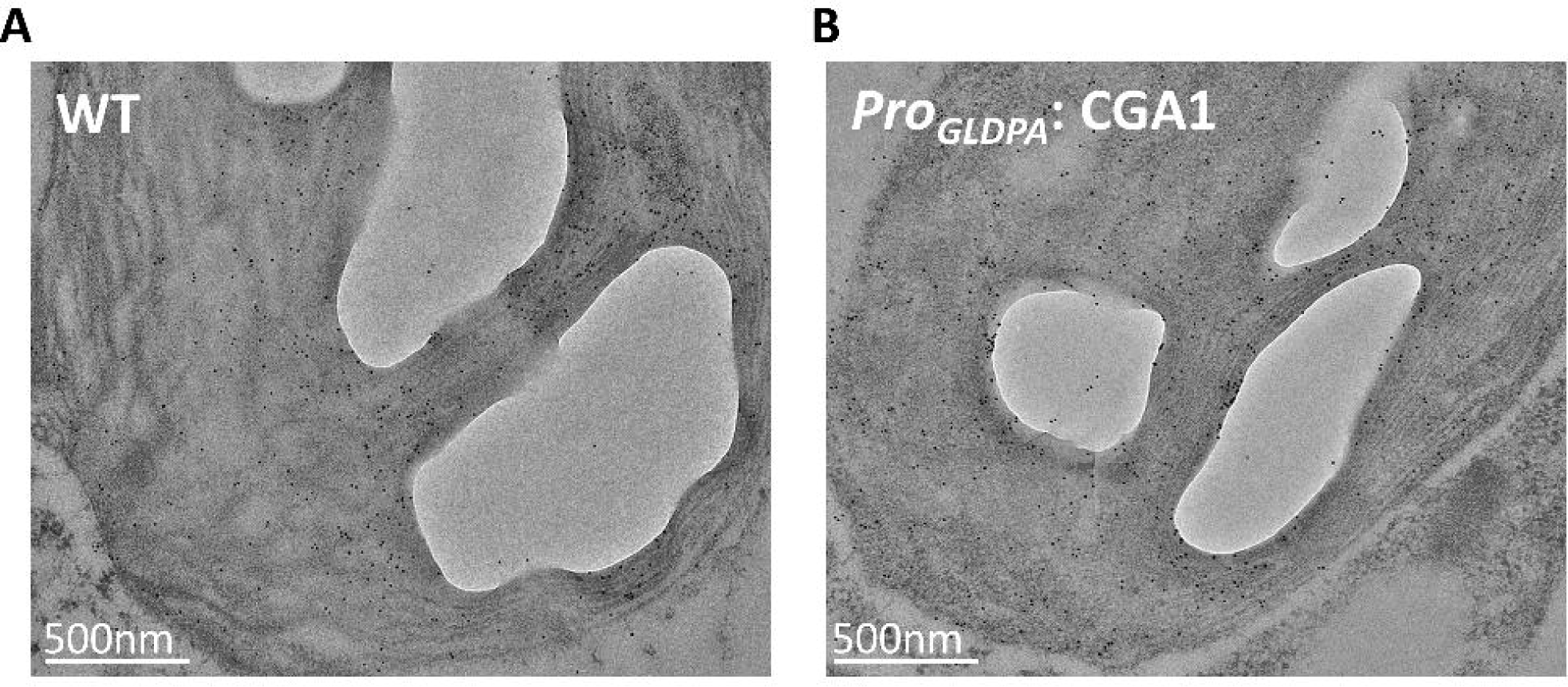
RuBisCO content examined with immunogold labelling. (A and B) Transmission electron micrographs of WT (A) and *Pro_GLDAP_*: CGA1 lines (B) showing plastid ultrastructure and immunogold labeling with RuBisCO antibody in BSC. Scale bars, 500 nm (A and B).

**Table 1.** Quantification of Rubisco in Bundle Sheath Cells. Mean ± SEM came from nine BSC from three individuals of WT and three transgenic *Pro_GLDAP_*: CGA1 lines (L3, L4, L9). *\*P < 0.05, **P < 0.01,***P<0.001*, Mann-Whitney U test.

|  | RuBisCO |  |  |
| --- | --- | --- | --- |
| | Total Gold<br>labeling / Cell | Density of Gold<br>labeling/Planar<br>chloroplast<br>Area ($\mu\text{m}^2$) | Density of Gold<br>Labeling/<br>Planar cell area<br>($\mu\text{m}^2$) |
| WT | 2986.0±422.0 | 106.9±8.7 | 11.8±1.0 |
| L3 | 5516.0±486.6** | 94.1±5.0 | 19.6±2.7* |
| L4 | 5251.0±570.4* | 107.8±7.1 | 18.9±1.8* |
| L9 | 6983.0±756.7*** | 93.0±5.7 | 18.2±1.7** |

### Expression of *CGA1* increased the number of mitochondria, peroxisome and plasmodesma and decreases photorespiration in bundle sheath

In the *CGA1* transgenic lines, we observed not only an increase in chloroplast number, but also increase in mitochondria and peroxisome numbers. Compared to the WT, the number of mitochondria in transgenic lines was 1.6 times higher (Fig 4A), and the number of peroxisomes was increased by about 1 fold (Fig 4C). Notably, the size of peroxisome in *Pro_GLDPA_*: CGA1 transgenic line showed inconsistent changes, since we observed an increase in L3 while a decrease in L9 (Fig 4D). The results indicate that *CGA1* may function not only in chloroplast development, but also in the development of other organelles. Compared with WT, more mitochondria located close to chloroplasts (Supplementary Fig 3C, D), which might enhance the refixation of photorespiratory CO_2_. In addition, the plasmodesma numbers between MC and BSC, between BSC and BSC as well as between BSC and VBC were also increased (Supplementary Fig 4), which strengthened the exchange and transport of metabolites between them.

**Fig 4.**
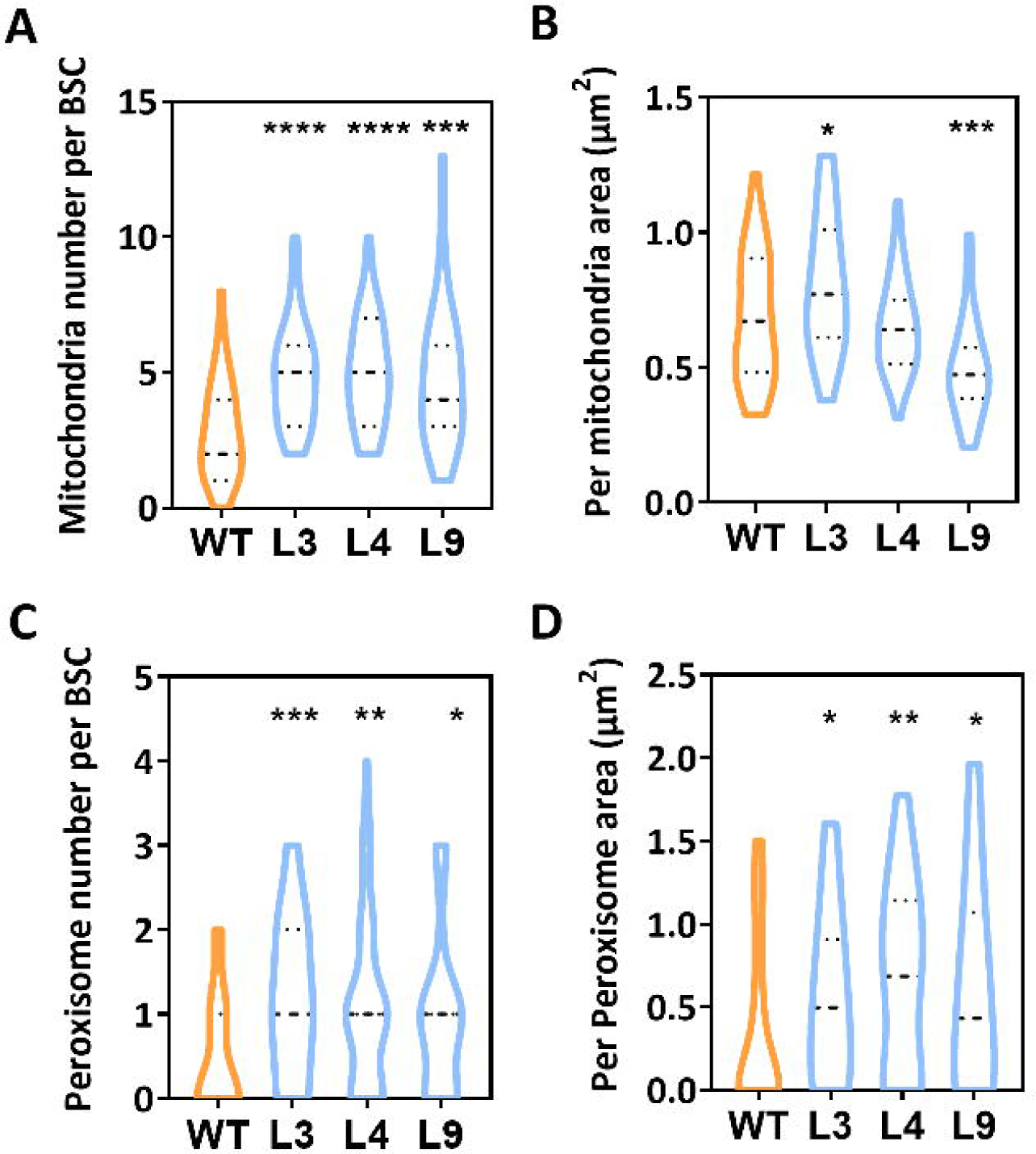
Expression of *CGA1* in the bundle sheath and vascular bundle increased mitochondria and peroxisome number and size. (A) The average mitochondria number in each BSC (n ≥39). (B) The average size of mitochondria in each BSC (n ≥39). (C) The average peroxisome number in each BSC (n ≥39). (B) The average size of peroxisome in each BSC (n ≥39*). *P < 0.05, **P < 0.01,***P<0.001*, two-sided student’s t-test, unpaired.

The glycine decarboxylase P-protein (GLDP), a major component of the glycine decarboxylase, is restricted in the BSC, which is viewed as a cellular feature for the intermediate photosynthetic type during the C_4_ evolution (Sage 2012; Schulze et al. 2016). We found that the density of immunogold labelling for GLDP was higher in the transgenic line than that in WT (Fig 5A, B). Specifically, the total immunogold labelling in BSC and the density of immunogold labelling in BSC area (um^2^) were increased in three independent transgenic lines compared with WT, but two lines showed significant differences (Table 2). The density of immunogold labelling in planar mitochondria area (μm^2^) were higher in the transgenic lines, although there is no significant difference in L9 and L4 (Table 2). These results indicated the abundance of GLDP was higher in the BSC in the transgenic line compared with WT, which was mainly due to the increased number of mitochondria.

**Fig 5.**
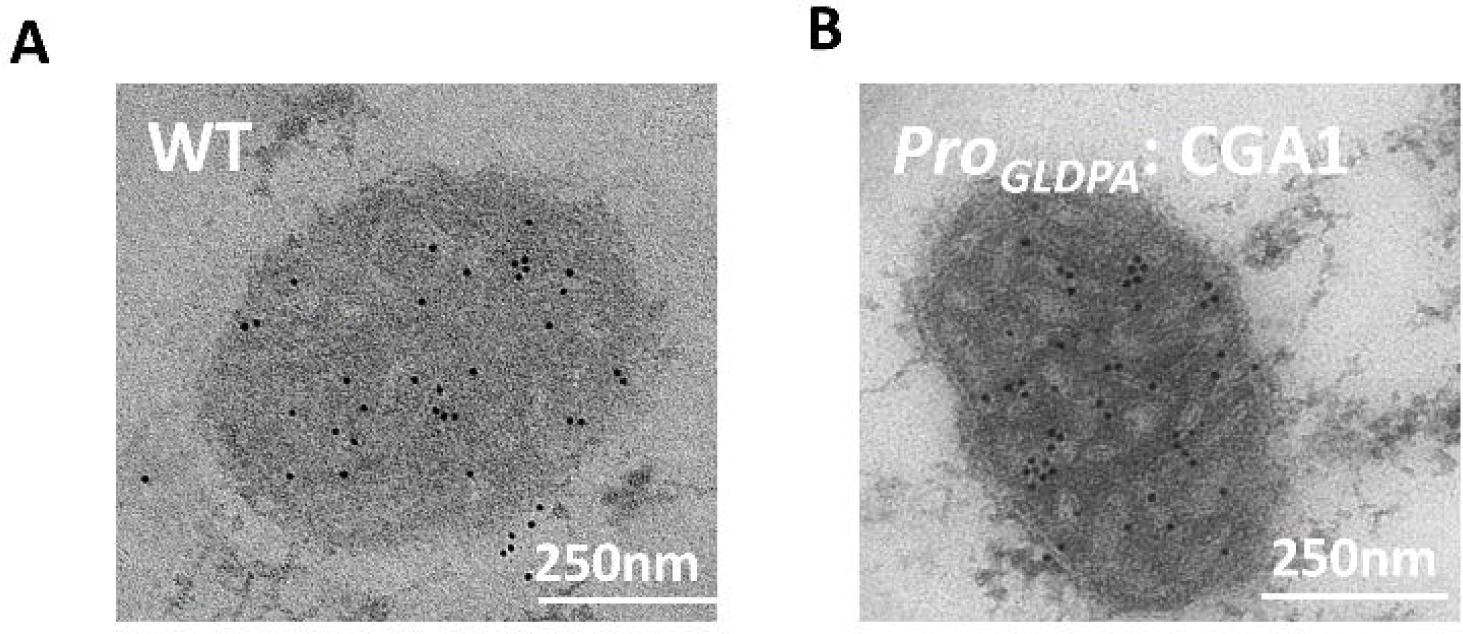
GLDP content examined with immunogold labeling. (E to F) Transmission electron micrographs of WT (E) and *Pro_GLDAP_*: CGA1 lines (F) showing mitochondria ultrastructure and immunogold labeling with GLDP antibody in BSC. Scale bars, 250 nm (E and F).

**Table 2.** Photosynthetic CO_2_ compensation points and day respiration were measured in WT and transgenic lines. Data are means of six biological replicates ± SD.

| Species | CO<sub>2</sub> compensation Point (ppm) | day respiration ($\mu\text{mol m}^{-2} \text{s}^{-1}$) |
| --- | --- | --- |
| WT | 51.2±1.14 | 0.99±0.13 |
| L3 | 49±0.8 * | 1.09±0.12 |
| L4 | 49.5±1.6 * | 1.06±0.14 |
| L9 | 49.4±1.3 * | 1.12±0.15 |

CO<sub>2</sub> compensation point was determined at 22°, 21% O<sub>2</sub>, and 250 $\mu\text{mol quanta m}^{-2} \text{s}^{-1}$ , two-sided student's t-test, unpaired.
Day respiration was determined by Light-response curve of leaf photosynthetic CO<sub>2</sub> uptake rate.

In addition, CO_2_ compensation points (Г), which is an indicator of the photorespiratory CO_2_ loss (Ku et al. 1991b), of three transgenic lines were significantly lower than that of WT (Table 3). Since we found no significant difference in R_d_ between transgenic lines and WT (Table 3), the decreased Г was due to reduced photorespiratory CO_2_ loss in transgenic line.

**Table 3.** Quantification of Glycine decarboxylase P-protein (GLDP) in Bundle Sheath Cells. Mean ± SEM came from nine BSC from three individuals of WT and three transgenic *Pro_GLDAP_*: CGA1 lines (L3, L4, L9). *\*P < 0.05, **P < 0.01*, Mann-Whitney U test.

|  | GLDP |  |  |
| --- | --- | --- | --- |
| | Total Glod labeling / Cell | Density of Gold labeling/Planar mitochondria Area ($\mu\text{m}^{-2}$) | Density of Glod Labeling/ Planar cell area ($\mu\text{m}^{-2}$) |
| WT | 121.8±18.8 | 127.1±7.5 | 0.4±0.1 |
| L3 | 247.1±26.0** | 195.6±17.5* | 0.8±0.1* |
| L4 | 191.1±31.7 | 149.2±19.0 | 0.9±0.1* |
| L9 | 234.9±33.4* | 158.3±11.2 | 0.6±0.1 |

### Expressing CGA1 in the bundle sheath and vascular sheath promotes increased leaf photosynthetic capacity

Consistent with the change in leaf anatomical features (Fig 2), we noticed increased photosynthetic rate (*A*) in transgenic lines under different CO_2_ and light levels compared to WT (Fig 6A, B). In addition, transgenic lines displayed higher stomatal conductance (*g_s_*) under different PPFD as well (Fig 6D). Therefore, the increased assimilatory CO_2_ was at least partially attributed to increased stomata conductance in the *Pro_GLDP_:CGA1*. Further analysis showed that there were higher maximum rate of RuBisCO carboxylation (V_cmax_) (Fig 6E) and higher maximum rate of electron transport (J_max_) (Fig 6F). Transgenic lines also showed increased amounts of large subunits of RuBisCO compared with WT (Supplementary Fig 5C). The percentage of vein area (VA) among total leaf area (TA) (Buhler et al. 2015) was similar to percentage of increase in photosynthetic rate (IP) among total photosynthetic rate (TP) (Fig 6C). Together, these results suggested that promoting the development of chloroplast in vascular bundle and bundle sheath improved leaf photosynthetic performance.

**Fig 6.**
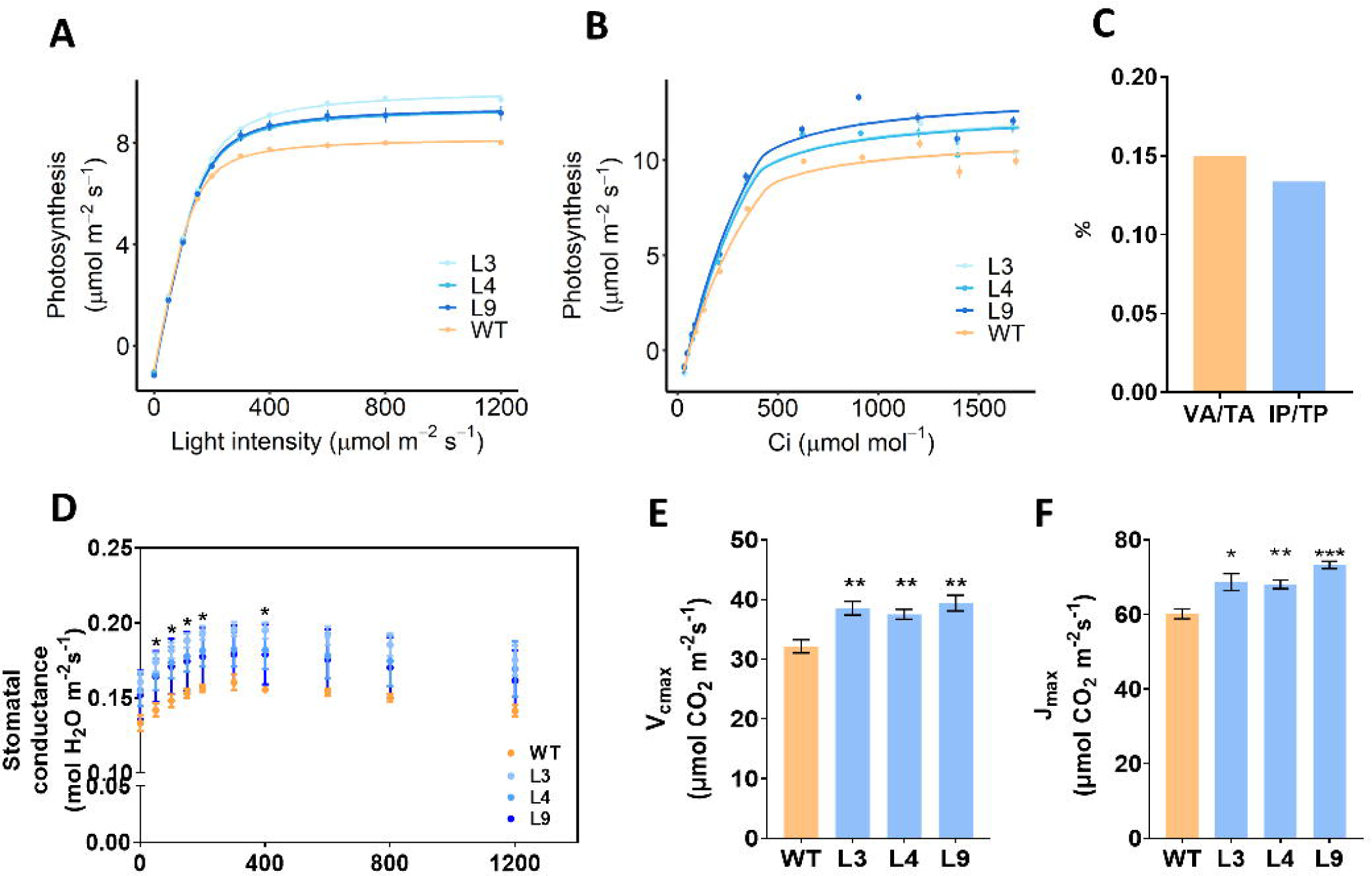
Expressing *CGA1* in the bundle sheath and vascular bundle promotes leaf photosynthetic capacity. (A) Light-response curve of leaf photosynthetic CO_2_ uptake rate (A-Q curve) fitted by a non-rectangular hyperbola model and measured at 22 □ and 400 ppm CO_2_ concentration. Data are presented as mean ± SEM (n=6). (B) CO_2_ response curve of leaf photosynthetic CO_2_ uptake rate (A-C_i_ curve) fitted by FvCB model and measured at 600 μmol m^−2^s^−1^ light intensity and 22□. Data are presented as mean ± SEM (n=4). (C) The comparison between the area of leaf veins occupies the area of the whole leaf and the percentage of increased photosynthesis and total leaf photosynthesis. VA/TA: The percentage between vein area and leaf area. IP/TP:The percentage between the increased part of photosynthetic rate and leaf total photosynthetic rate. (D) Light-response of stomatal conductance measured at 22 □ and 400 ppm CO_2_ concentration. Data are presented as mean ± SEM (n=6). *\*p<0.05* for L3, L4, and L9 compared with WT, Student’s *t-*test. (E and F) Maximum rates of carboxylation (V_cmax_) (E) electron transport (J_max_) (F) in WT and L3, L4, and L9. Values were estimated by the fitted FvCB model. Data are presented as mean ± SEM (n=4). All measurements in (A) to (F) were performed with rosette leaves of plants growth in phytotron at the 28-day-old. \**P* < 0.05, \*\**P* < 0.01,***P<0.001, Student’s *t*-test.

### Expression of CGA1 in the vascular bundle and bundle sheath enhanced biomass production

The biomass under either normal or high light intensity were examined. Compared with WT, transgenic lines exhibited denser canopies under either normal (Fig 7A) or high light intensity (Fig 7 D). The biomass and the leaf number in the *Pro_GLDPA_: CGA1* were higher than those in WT under either normal light (Fig7 B, C) or high light levels (Fig 7E, F). The increase in biomass and leaf number of transgenic lines under high light intensity were higher than that under low light intensity (Fig 7B, C, E, F), which was consistent with the higher increase in photosynthetic rate of transgenic lines under high light intensity compared to low light intensity (Fig 6A).

**Fig 7.**
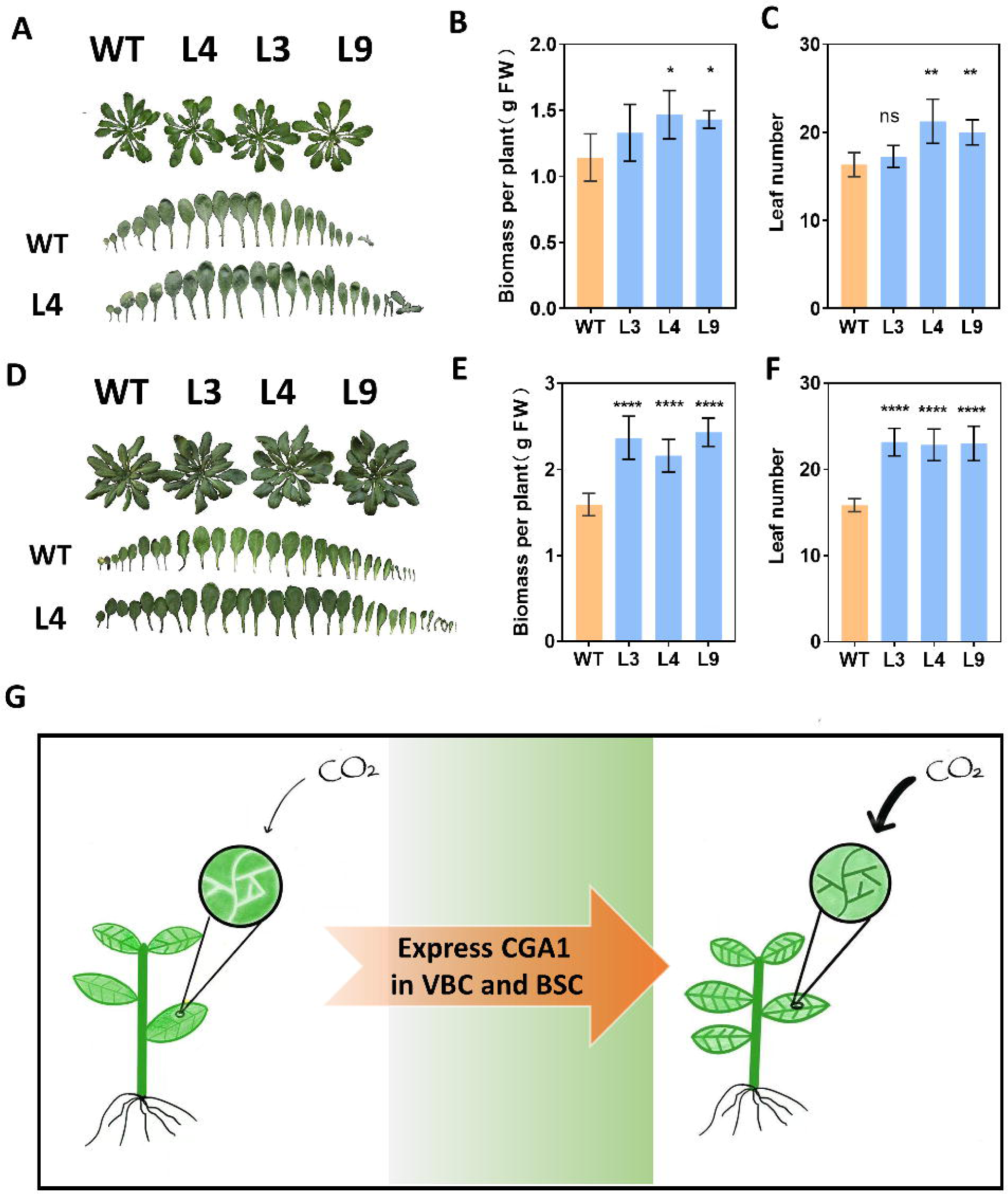
Expression of *CGA1* in the bundle sheath and vascular bundle increases biomass and leaf number. The photograph of phenotype under normal light (A) and high light (D). The biomass and leaf number of plants grown under normal light (B, C) and high light (E, F). (G) Schematic diagram of expressing *CGA1* in the BSC and VBC. Expression of *CGA1* in the BSC and VBC activates BSC and VBC which primarily lacks photosynthesis machinery. The activated BSC improves photosynthetic CO_2_ uptake of leaves and increased whole plant photosynthesis and biomass production. \**P* < 0.05, \*\**P* < 0.01, ***P<0.001, Student’s *t*-test.

## Discussion

### CGA1 can be used to activate photosynthesis in vascular bundle and bundle sheath cell

This work indicates that the photosynthetic activity of the bundle sheath cell (BSC) and vascular bundle cell (VBC) can be increased by expression of *CGA1* in these cell types. First, the starch granule number was increased in VBC and BSC in *Pro_GLDPA_*: *CGA1* relative to WT (Fig 2H, N). Secondly, the content of RuBisCO was increased in the *Pro_GLDPA_*: *CGA1* relative to WT (Fig 3, table1). Thirdly, the leaf photosynthetic rate was largely increased in the *Pro_GLDPA_*: *CGA1* compared with WT (Fig 6A, B). We found no expression in mesophyll cell (MC) under the controlling of *GLDP* promoter from *Flaveria trinervia* (Engelmann et al. 2008)(Supplementary fig 1). Since *CGA1* was specifically expressed in VBC and BSC (Engelmann et al. 2008) (Supplementary Fig 1), and the chloroplasts of MC didn’t show difference between WT and *Pro_GLDPA_*: *CGA1* (Suppmentary fig 3), suggesting the increased photosynthetic rate was attribute to increased photosynthesis in BSC and VBC. Consistent with this, earlier studies have also shown that expression of *GLK* in the root of *Arabidopsis* leads to fully functional photosynthetic tissues in *Arabidopsis* root (Koichi Kobayashi and Masuda 2013). Therefore, expression of transcription factors which can promote the development of chloroplasts (Cackett et al. 2022; Yi-Hsuan Chiang 2012; Zubo et al. 2018) represents a promising approach to improve photosynthesis in photosynthetically inactive cells. Considering that these controller for chloroplast development have been shown to be able to function in almost all plant tissues examined (Koichi Kobayashi and Masuda 2013; Wang et al. 2017; Yi-Hsuan Chiang 2012; Nakamura et al. 2009), the potential to improve photosynthetic capacity in photosynthetically inactive cells through activating chloroplast development could be explored in more plant tissues.

Here we would like to mention that *CGA1* and *GLK* have been expressed in BSC and VBC of rice, but no enhancement in photosynthetic performance was reported (Dong-Yeon Lee 2021; Wang et al. 2017; Nomura et al. 2005; Engelmann et al. 2008). One reason for this difference between these earlier studies and current one might be due to the differential degree of enhancement on chloroplast development. Both the chloroplast number and size were significantly increased in the BSC in this study (Fig 2I, J), but only the chloroplast size was significantly increased in previous work (Wang et al. 2017; Dong-Yeon Lee 2021). Our results also showed that, compared to plant performance under lower light intensity, there was a larger difference in photosynthesis and biomass accumulation under higher light intensity between *Pro_GLDPA_*: *CGA1* and WT (Fig 3A; Fig 6E), which indicated that in the earlier studies, the light level used in the experiments might be not sufficient to induce the expected increase in photosynthesis (Wang et al. 2017; Dong-Yeon Lee 2021). Under low light intensity, the amount of light reaching BSC might be low, which compromised the ability of BSC to capitalize on the photosynthesis.

### Activating photosynthetic capacity of vascular bundle and bundle sheath cell in C_3_ leaves improved leaf photosynthesis

Different strategies to improve the plant photosynthesis to have been mainly related to improve photosynthetic capacity of the MC (South et al. 2019; De Souza et al. 2022). This work indicated that improving photosynthetic capacity of BSC and VBC by specific expression of *CGA1* in these tissues can substantially improve leaf photosynthesis as well (Fig 6), resulting in the improvement of whole plant biomass (Fig 7). Interestingly, the percentage of leaf vein area relative to total leaf area was almost equal to the percentage increase of photosynthesis in *Pro_GLDPA_:CGA1* relative to WT (Fig 6C), suggesting that photosynthetic capacity of bundle sheath and vascular bundle in the transgenic plants is similar to that of mesophyll (Fig 5A, B). Our study supported the notion that the cell deficient in photosynthetic capacity has the larger potential to be used to improve the entire plant photosynthesis.

The enhanced photosynthesis in the *Pro_GLDPA_:CGA1* might be related to the increased refixation of photorespiratory release of CO_2_. During C_4_ evolution, the ability to recapture CO_2_ released from photorespiration in the proto-Kranz has been considered to be enhanced (Schluter and Weber 2016; Sage et al. 2013; Ku et al. 1991a). Compared with WT, CO_2_ compensation point (Г), which is related to refixation of photorespiratory release of CO_2_ (Ku et al. 1991a; Schluter and Weber 2016; Sage et al. 2013), was decreased in *Pro_GLDPA_: CGA1* (Table 3), as the species with proto-Kranz (Sage et al. 2013). Theoretically, Г can be potentially influenced by respiration and photorespiration (Sage et al. 2013; Ku et al. 1991a). Here we found no difference in respiration between *Pro_GLDPA_*: CGA1 and WT (Table 3), indicating the apparent change in Г could be attributed to changes in photorespiration. In the *Pro_GLDPA_:CGA1* transgenic plants, the development of chloroplast and mitochondria were all enhanced (Fig 2, Fig 4), and BSC in *Pro_GLDPA_:CGA1* had the higher GLDP content (Table 2), together resembling the feature of proto-Kranz; furthermore, mitochondria was closer to the interior than chloroplasts in BSC (Supplementary fig 3), and the VBC and BSC were located in the interior of leaves; all these might increase the percentage of refixation of photorespired CO_2_.

### CGA1 being a major molecular tool used to mimic the step of proto-Kranz formation during C_4_ evolution

Engineering of C_4_ photosynthesis into the C_3_ plants have long been considered as an effective approach to increase the photosynthetic capacity of C_3_ plants (Wang et al. 2017; Schuler et al. 2016). The most recent advance on this is the installation of C_4_ biochemical pathways into rice (Ermakova et al. 2021). However, the installed pathway did not show expected C_4_ flux (Ermakova et al. 2021). Concurrent with the direct installation of C_4_ pathways into C_3_ plants, another strategy to create C_4_ photosynthesis is to mimic the process of C_4_ evolution (Heckmann et al. 2013), i.e. to repeat the process of proto-Kranz, C_2_, C_4_-like species sequentially from C_3_ plants (Sage et al. 2014; Sage et al. 2012; Zhao et al. 2022). According to the current pyramid theory of C_4_ evolution, different anatomical, biochemical and physiological traits were acquired during the different steps of the C_4_ evolution (Sage et al. 2012). This study suggested that expression of *CGA1* in BSC and VBC enabled *Arabidopsis* to acquire many anatomical, biochemical and physiological features of a plant with proto-Kranz (Fig 3, Fig5, table 1) (Sage et al. 2013). The enhanced development of chloroplast, mitochondria and peroxisome, increased plasmodesmatal connection, the increased starch granule number, increased content of RuBisCO and GLDP, and changed photosynthetic parameters (photosynthetic rate, CO_2_ compensation point) (Fig 3, table 1, Fig 5, table 2, Fig 2H, Fig 6A,B) all suggested that the engineered *Arabidopsis* acquired the key features for a leaf with initial C_4_ (Wang et al. 2017; Sage 2012). Besides being an intermediate stage during the evolution of C_4_ photosynthesis from C_3_ photosynthesis, plants with C_2_ photosynthesis could have the advantage in the conditions with the high potential of photorespiration, such as hot and drought (Sage et al. 2012; Lundgren 2020). Whether *Pro_GLDPA_:CGA1* would gain increased stress resistance still needs to be tested.

In addition to the earlier observation that expression of *CGA1 or GLK* could induce development of chloroplast and mitochondria (Dong-Yeon Lee 2021; Wang et al. 2017), here also found that peroxisome development was induced in BSC when *CGA1* was expressed in BSC (Fig 4C, D). The development of these organelles in BSC is highly coordinated during C_4_ evolution and development (Sage et al. 2013; Stata et al. 2019; R. HAROLD BROWN 1983). Given the close coordination between organelles are critical for the normal function of any plant cell, including BSCs and VBCs (de Souza et al. 2017), identifying methods to enable coordinated developmental changes of these organelles is probably needed to ensure a successful engineering for C_3_ cells. Here results from this study suggested that a single transcription factor *CGA1* could be used to coordinate development of these three organelles in BSC, and hence might be involved in C_4_ evolution and development.

## Acknowledgments

This work is supported by Strategic Priority Research Program of the Chinese Academy of Sciences (Grant No. XDB27020105), National Research and Development Program of Ministry of Science and Technology of China (2020YFA0907600).

We thank Jilei Huang, Chuanhe Liu and Xiaoxian Wu from Instrumental Analysis & Research Center, South China Agricultural University for helping on TEM sample processing and image acquisition.

## Author Contributions

X.G.Z, F.F.M, Y.Y.Z designed the research; F.F.M, performed transgenic, electron microscopy and gas-exchanged experiment; Y.Y.Z and F.F.M took the photos of plants; Y.Y.Z and F.F.M plotted the graphs and explained the data; X.G.Z, Y.Y.Z, F.F.M wrote the paper; B.H.S performed the immunogold labeling experiments. F.S.L performed the model fitting; M.J.A..L help analyze the transcriptome.

## Notes

### Competing Interest Statement

The authors have declared no competing interest.

