## Supplementary figures and images for "Enhancing plant photosynthesis through activating cells deficient in photosynthetic apparatus"

### supplmentary fig 1

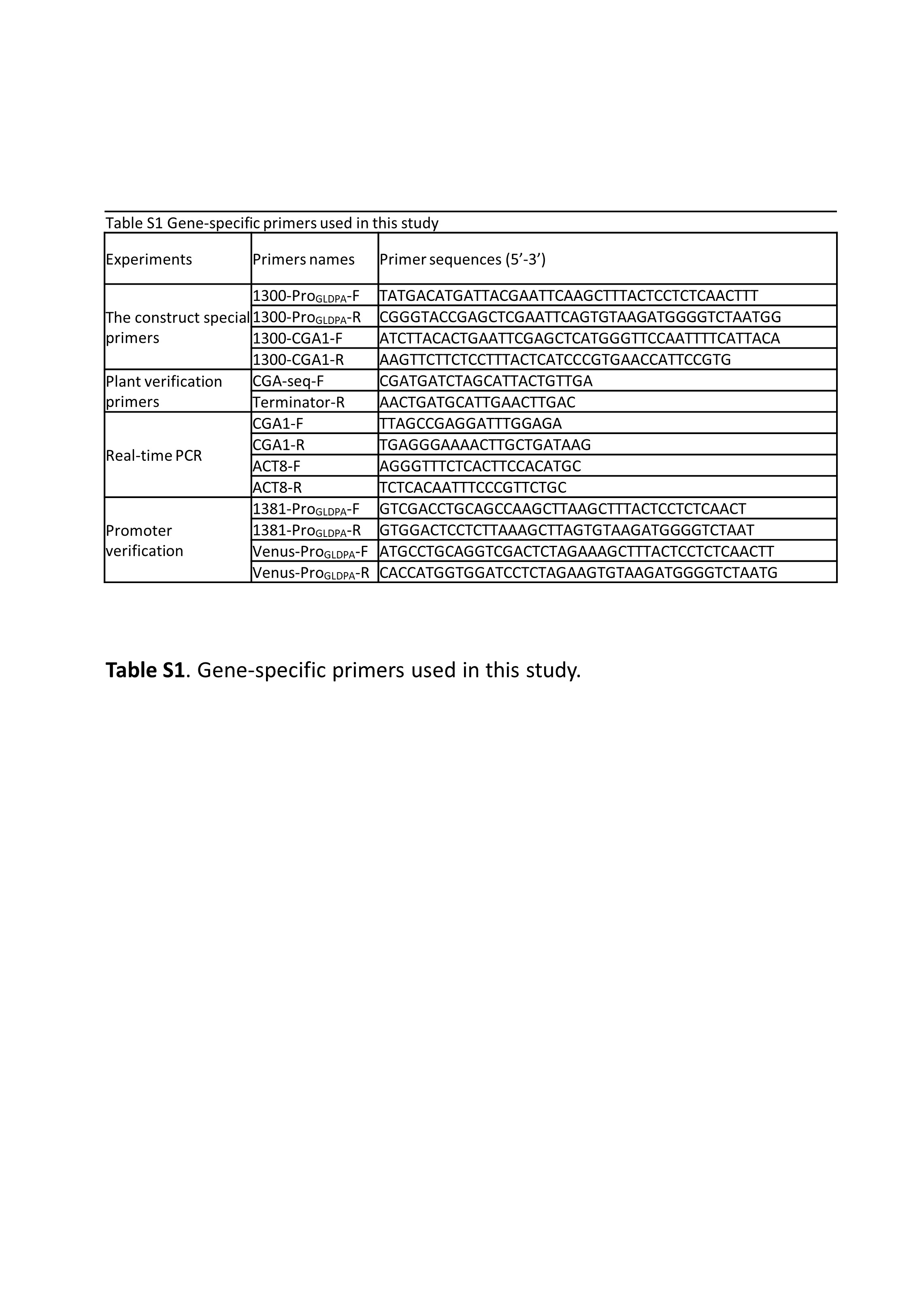

### supplmentary fig 2

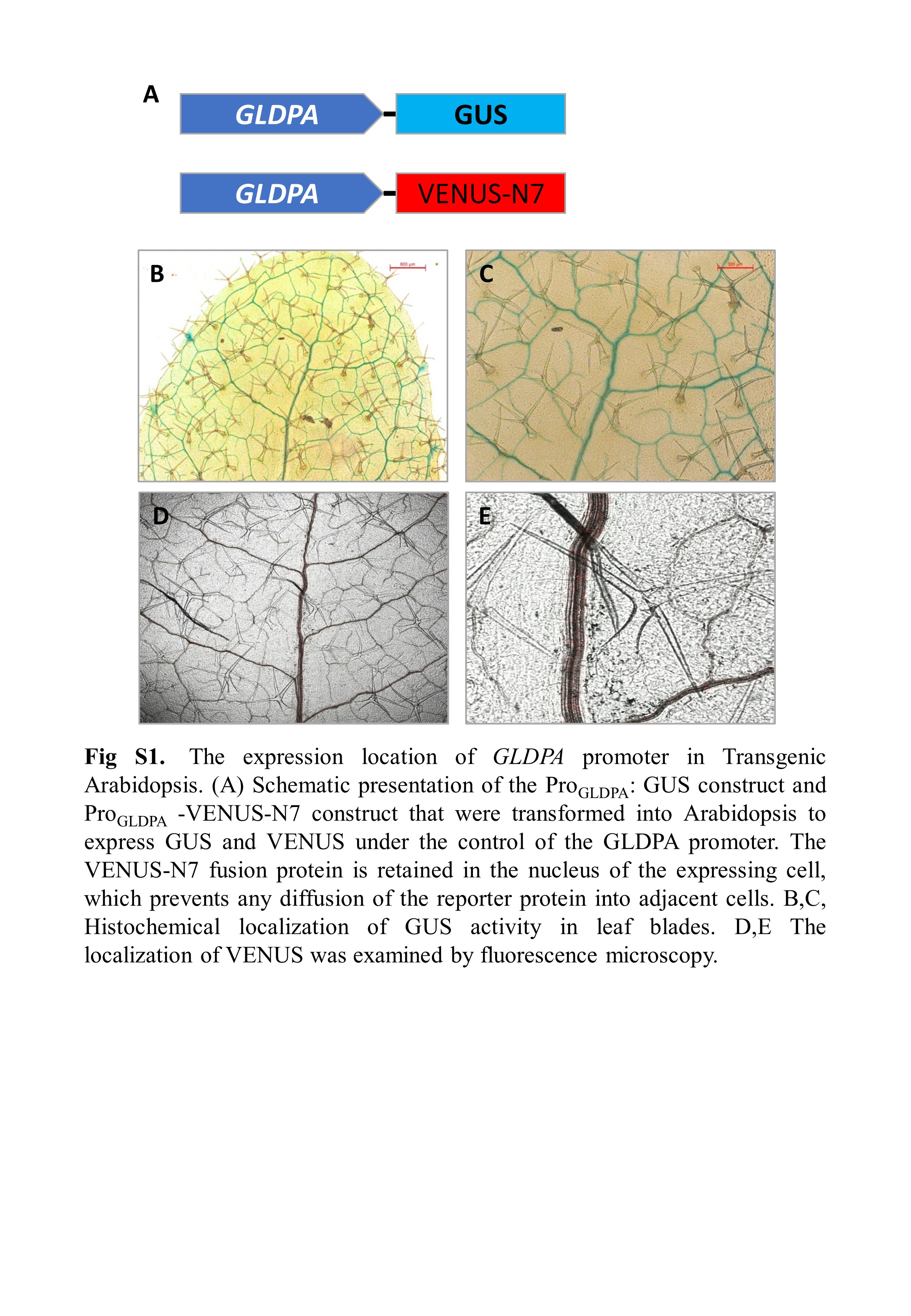

### supplmentary fig 3

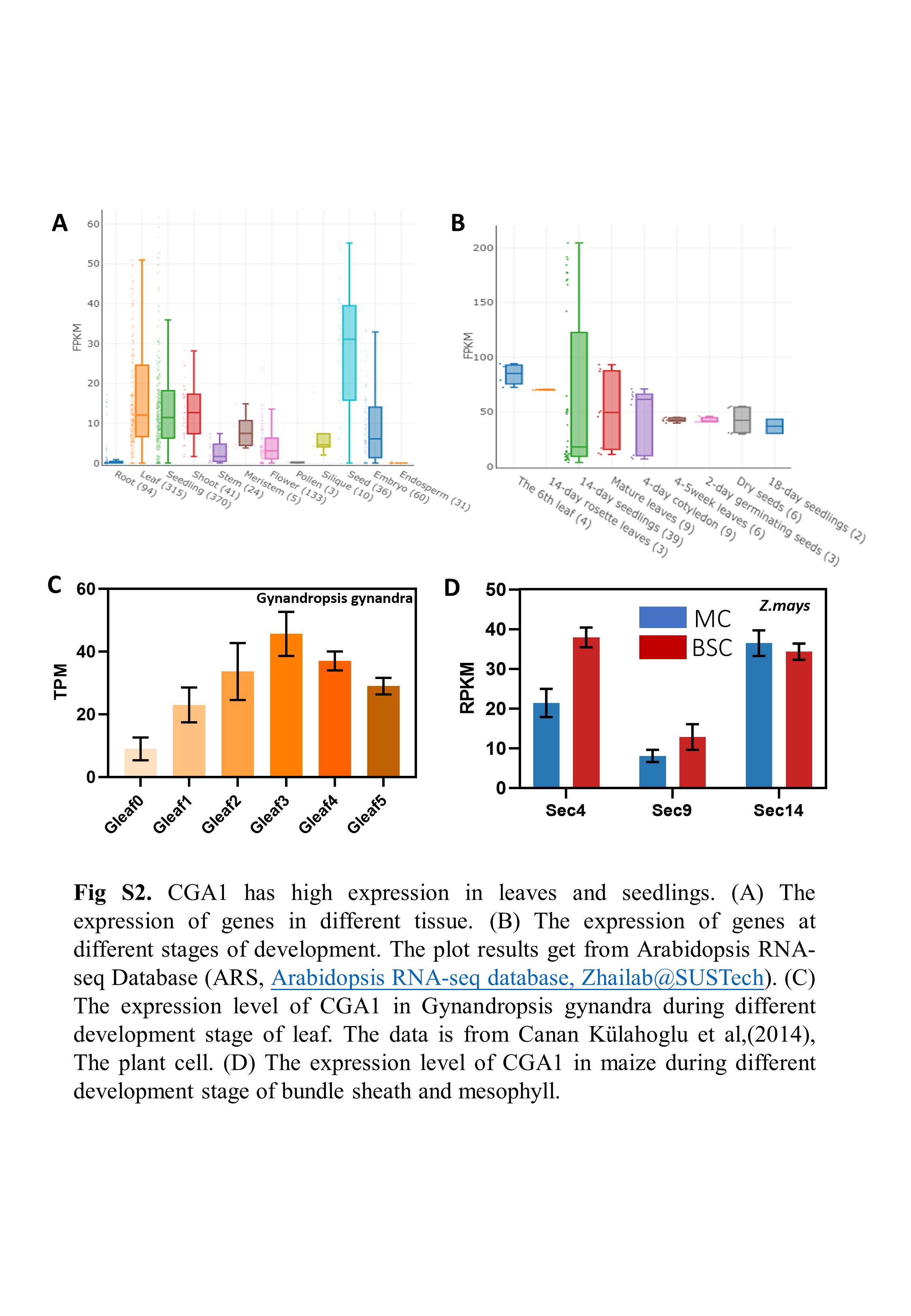

### supplmentary fig 4

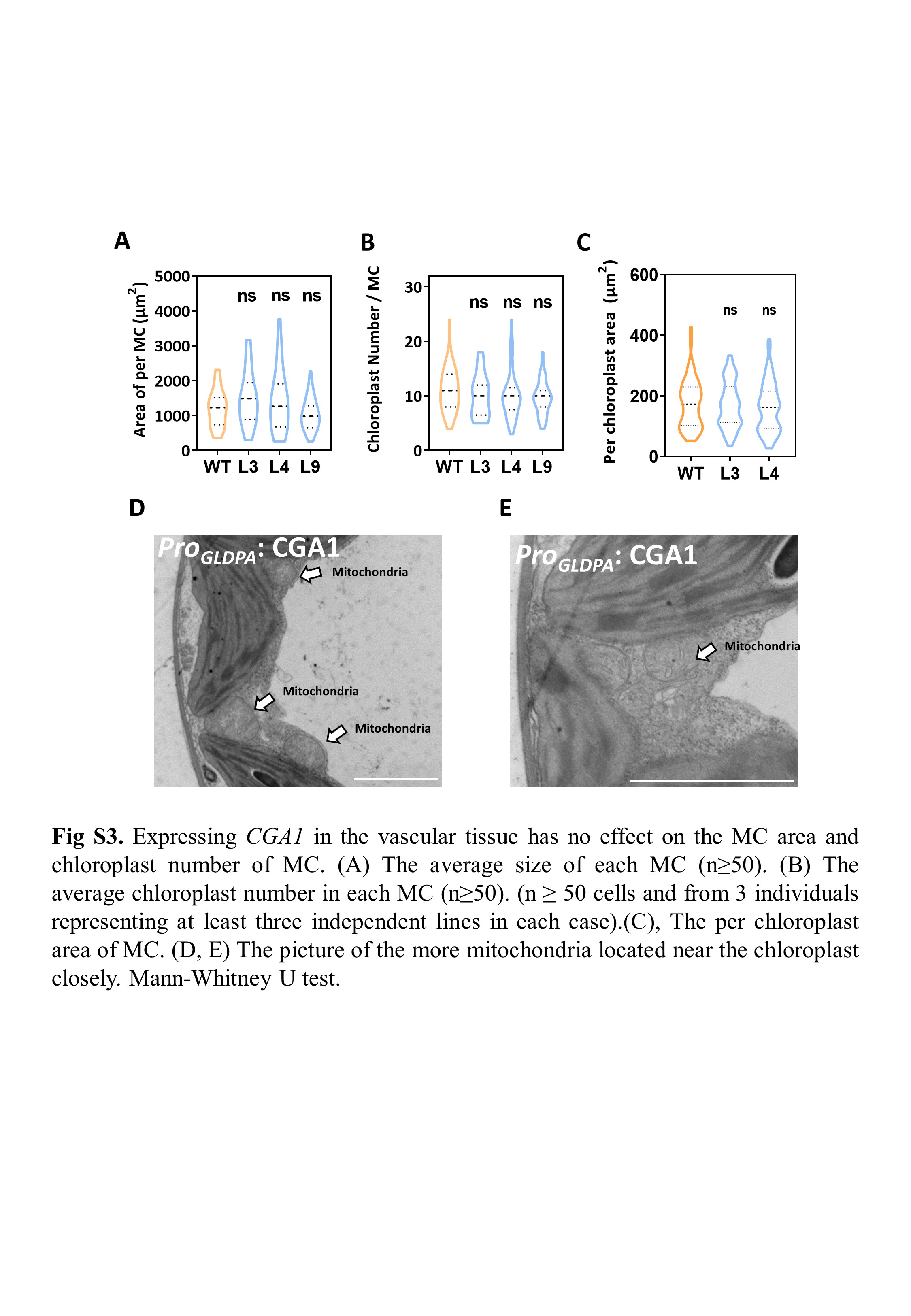

### supplmentary fig 5

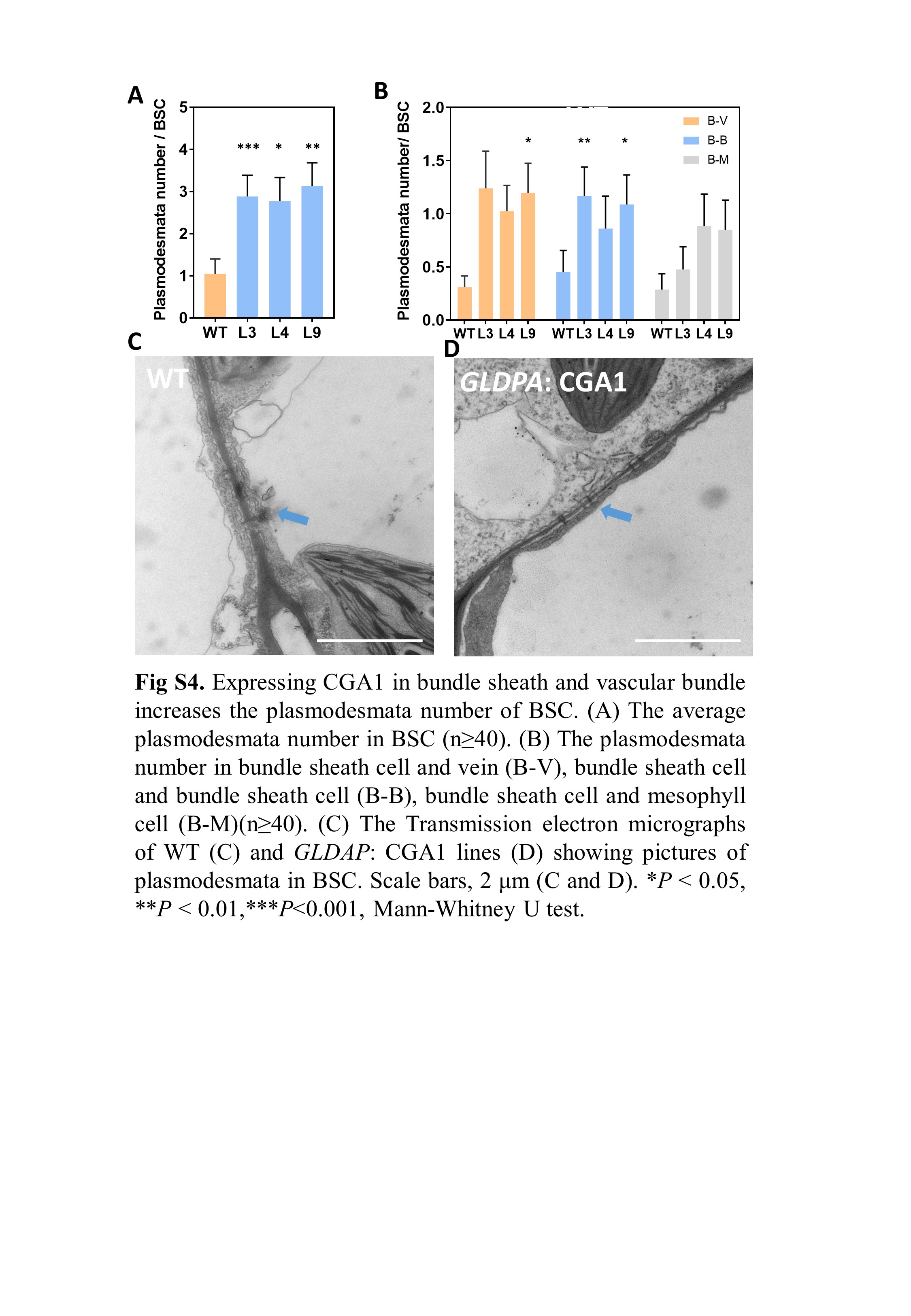

### supplmentary fig 6

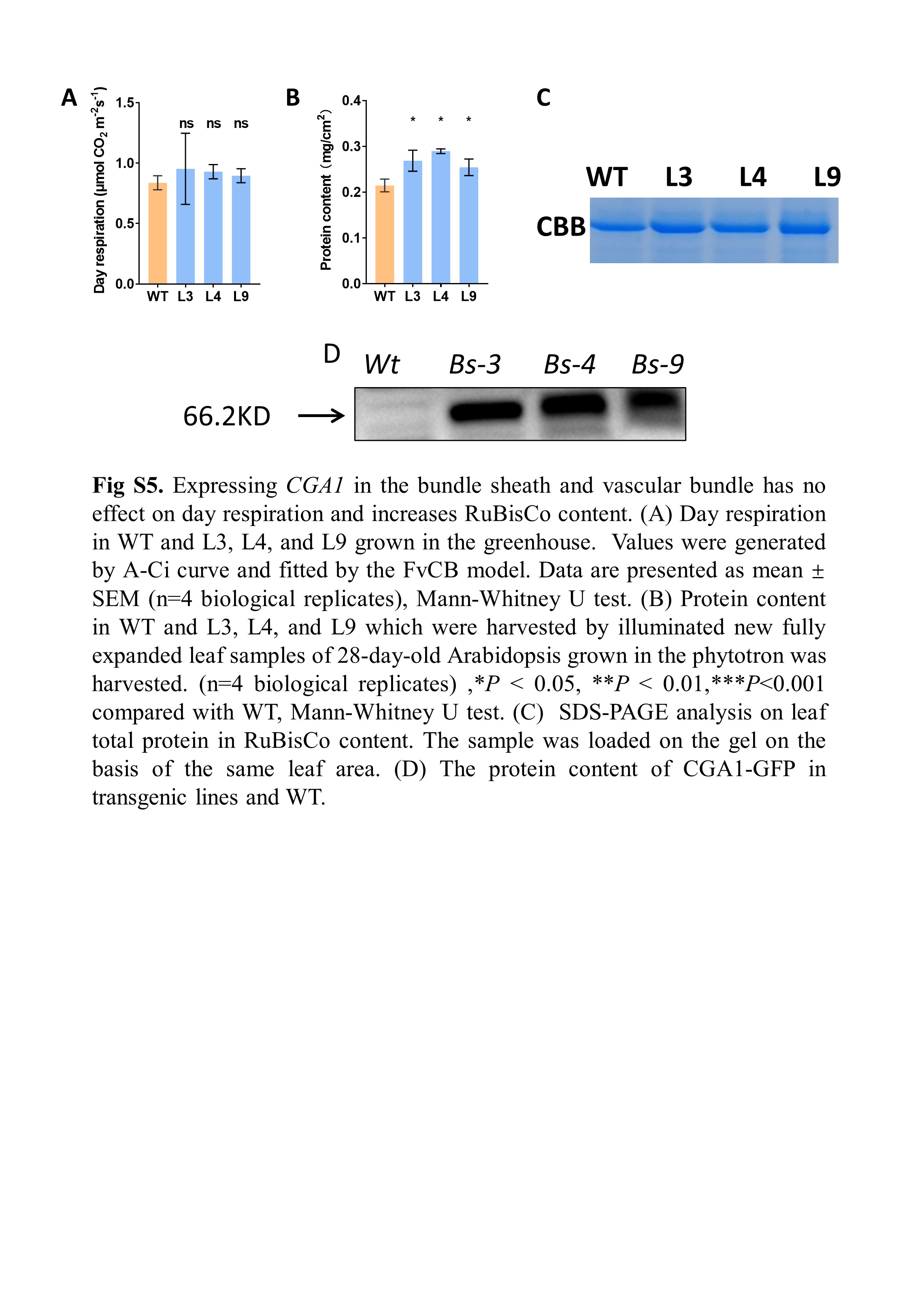

### supplmentary fig 7

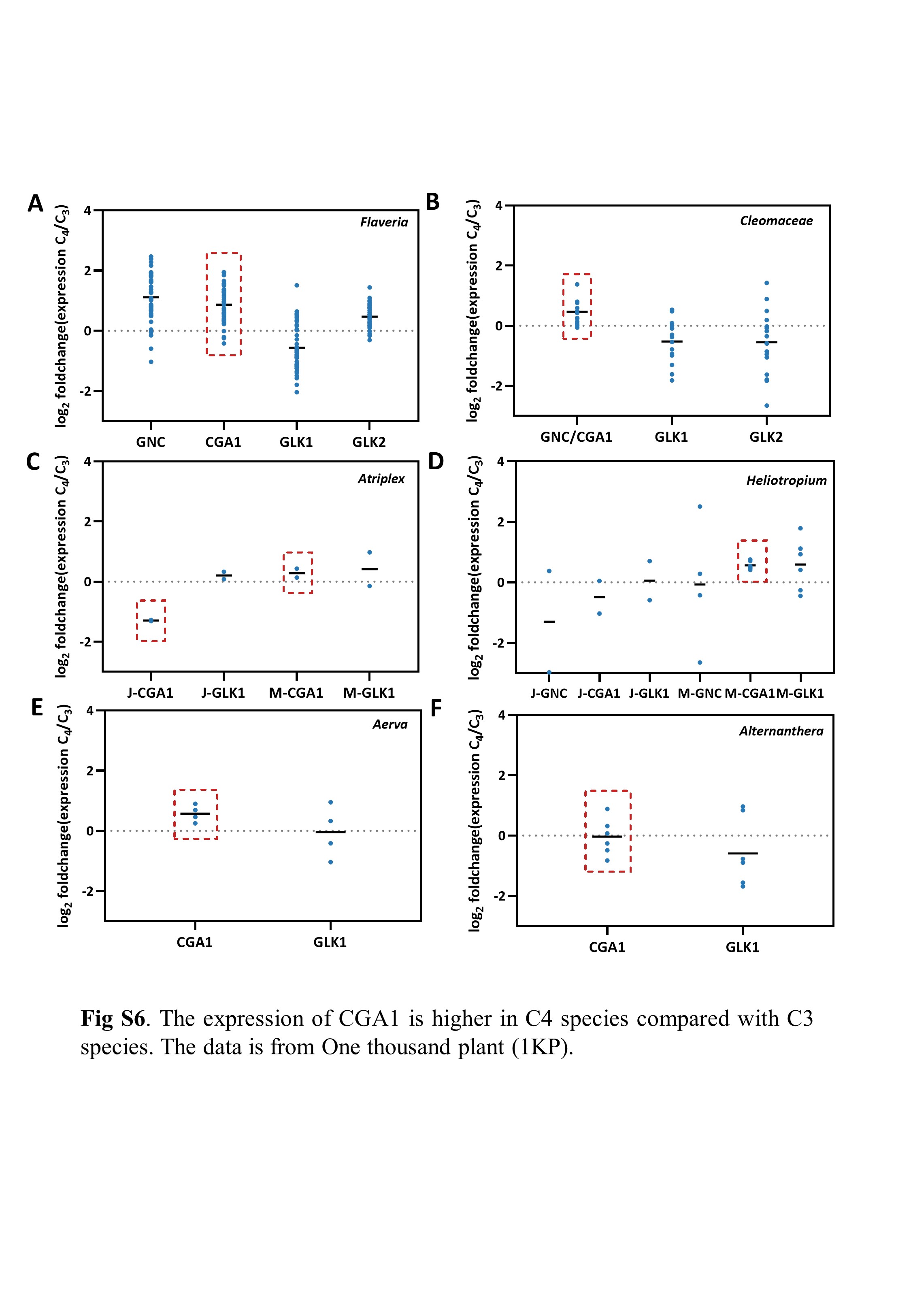
